# A Milk Fat Globule Membrane-enriched dairy co-product modulates gut *Faecalibaculum rodentium* metabolism in association with the prevention of cognitive impairment in aging male Wistar rats

**DOI:** 10.1101/2025.04.02.646113

**Authors:** Kasey Schalich, You-Tae Kim, Duncan Sylvestre, Aidong Wang, Gulustan Ozturk, Karen Kalanetra, Daniela Barile, Ameer Taha, David Mills

**Author notes:** To whom correspondence should be addressed. David Mills, Department of Food Science and Technology, College of Agriculture and Environmental Sciences, University of California Davis, One Shields Avenue, Davis, CA, USA 95616.

## Abstract

Consumption of the milk fat globule membrane (MFGM) by infants is linked to enhanced neurodevelopment and sustained cognitive improvements later in life. Aging, by contrast, is often marked by neurodegeneration and cognitive decline-a growing concern in the U.S. as 6.9 million Americans live with Alzheimer’s disease (AD). As such, identifying interventions to prevent cognitive impairment are imperative. The whey protein phospholipid concentrate (WPPC), a dairy co-product, is enriched in MFGM glycoconjugates. Given the cognitive health benefits conferred by MFGM consumption in early life, we previously found that high-fat (HF) diet induced cognitive impairment in aging male Wistar rats was prevented by supplementating with a 1.6% or 10% WPPC in the diet, compared to control rats fed a low-fat (LF) diet. We hypothesized that WPPC exerts protective effects against cognitive impairment through the gut-brain axis by modulating gut microbial composition and metabolism. To test this, we analyzed 16S rRNA sequencing data from fecal samples of aged male Wistar rat (4 months old) fed a LF, HF, HF + 1.6% WPPC, or HF + 10% WPPC diet (n=9-10/group). Compared to LF, the HF diet reduced the abundance of the Erysipelotrichaceae family, particularly the species *Faecalibaculum rodentium*, which increased numerically with the 10% WPPC diet. Interestingly, Erysipelotrichaceae relative abundance correlated with hippocampal memory storage (spearman correlation=0.4, p=0.034). In vitro growth assays confirmed that *F. rodentium* grew robustly in isolation on WPPC glycoconjugates and on constituent WPPC components (p<0.05). RNA sequencing of F*. rodentium* grown on the WPPC MFGM glycoconjugates versus glucose (n=3/group) revealed significant upregulation of genes involved in in amino acid metabolism and fatty acid oxidation (FDR<0.05). Collectively this data suggests a potential role for *F. rodentium* in preventing cognitive impairment through the gut-brain axis by metabolizing the WPPC that may act on the host.

## Introduction

With few exceptions, the ability to synthesize and secrete milk is unique to the class *Mammalia*. Milk is a complex biofluid which undergoes dynamic compositional variance over the course of lactation to be in synchrony with the evolving nutritional, growth and developmental needs of the offspring. Milk contains both nutritive and bioactive components that play crucial roles in promoting enteric and systemic development, while shaping the establishment and function of the nascent gut microbiome. The milk fat globule membrane (MFGM) is a structurally complex and bioactive component in milk, with species-specific compositional differences.^1^ During lactation, lipid droplets containing a triglyceride core undergo anterograde transport from the endoplasmic reticulum to the plasma membrane of mammary epithelial cells, where they are enveloped by the plasma membrane during excretion. This process forms the MFGM’s distinctive tri-layer structure, composed of approximately 22% proteins (predominantly glycoproteins) and 72% lipids (predominantly phospholipids). Notably, many MFGM proteins and lipids are glycoconjugates with diverse bioactive properties, including promoting the viability of probiotic bacteria,^2–4^ reducing inflammation and intestinal permeability,^5–7^ mitigating oxidative stress,^8^ accelerating gut and mucosal immune system maturation,^9,10^ increasing intestinal neurotransmitter abundance,^10^ alleviating stress-induced cognitive defects,^11^ and improving cognitive impairment,^12^ physical performance,^13^ and survival rates with aging.^5^

Accumulating evidence suggests that the MFGM actively contributes to neonatal neurodevelopment and is linked to enhanced cognitive function and neurodevelopmental outcomes across species, including rodents,^5,14,15^ piglets,^16^ and humans.^17–21^ In infants, supplementation of formula with MFGM-enriched fractions has been shown to modulate immune responses,^22^ shape gut microbiota composition and its associated metabolome,^23–25^ enhance short-term memory,^26^ and improve behavioral outcomes such as increased sleep and improved expressive language, social emotional and general adaptive behaviors.^21,26,27^ Intriguingly, MFGM consumption during infancy-whether through breast milk or formula-has been associated with long-term cognitive benefits, including improved school-age cognitive performance,^28,29^ suggesting its potential impact across the lifespan. More recently, studies in adults have demonstrated that MFGM supplementation enhances physical performance^13^ along with stress resilience and sleep quality,^30^ indicating that its neuroprotective and cognitive benefits extend into adulthood and late life.

In the industrialized world, longer lifespans enabled by societal advancements have been accompanied by rising rates of neurodegeneration and cognitive impairments.^31,32^ In 2024 in the United States, 5% of people aged 65-74 years old, 13.2% of people aged 75-84, and 33.4% of people aged 85 years or older have Alzheimer’s dementia.^33^ Approximately 50% of individuals > 95 years old have Alzheimer’s disease.^34^ Beyond genetic predisposition, modifiable risks factors such as elevated body mass index (BMI)-often driven by the Western diet, characterized by high-fat and high-sugar intake-appear to contribute to neurodegenerative risk.^33,34^ Interestingly, according to a dementia prevention report by the Lancet Commission, it is estimated that 40% of dementia cases can be attributed to modifiable risk factors, suggesting that nutritional interventions may mitigate disease prevelance.^33,34^

Recognizing the crucial role of MFGM in supporting neuronal development and function during early life, we explored whether MFGM could offer similar protective effects against neurodegeneration and cognitive decline in later life. Our previous work demonstrated through behavioral assessments and measurement of the evoked response that supplementing aging male Wistar rats on a high-fat (HF) diet with a MFGM-enriched dairy fraction -the whey protein phospholipid concentrate, or WPPC-significantly preserved working memory function compared to a low fat (LF) diet control group (Sylvestre, et al. 2025, on bioRxiv). These findings align with prior research showing that MFGM-enriched supplementation can mitigate cognitive decline and neurodegeneration in rodent models.^12^ Given the abundance and structural complexity of the MFGM glycoconjugates in the WPPC, we hypothesized that the WPPC’s neuroprotective effects might be mediated through modulation of gut microbiota function, thus engaging the gut-brain axis as a mechanism of action. As such, this study aimed to characterize how WPPC consumption influences gut microbiome composition in aging male Wistar rats, and to identify specific microbial metabolic shifts associated with improved cognitive outcomes. As such, this work provides insight into potential nutrient-gut microbe-brain interactions underlying MFGM’s (via WPPC) protective affects against cognitive impairment.

## Materials and Methods

### Collection of rat colon content

Colon content was collected between 11am and 6pm from rats in our previously published study (Ref Duncan’s paper). From each experimental group (LF diet, HF diet, HF+1.6% WPPC, and HF+10% WPPC), feces was collected directly from the colon of the individual rats following sacrifice. Colon content was immediately frozen on dry ice and then stored at −80°C until further use.

### 16S rRNA sequencing of rat colon content

Genomic DNA was extracted with Zymobiomics DNA Miniprep Kit (Zymo Research, Irvine, CA) following the manufacturer’s instructions. The resulting DNA was PCR amplified in triplicate with barcoded primers targeting the 16S rRNA gene as described in the previous study (https://doi.org/10.1371/journal.pone.0066437). The raw sequencing reads, fastq files, were demultiplexed with Sabre and imported into QIIME2-2022.11 (https://doi.org/10.1038/s41587-019-0209-9). After filtering, taxonomy was assigned to reads using a pre-generated naïve Bayes classifier trained on Silva database version 138 (99% identity) amplicon sequence variants (ASVs) from the 515F/806R region of sequences (https://doi.org/10.1186/s40168-018-0470-z, https://doi.org/10.1093/nar/gks1219, https://doi.org/10.1038/ismej.2012.8). All samples were normalized to 33864, the smallest read number among the samples. Additional analyses and visualization were perfumed using R 4.4 statistical software with packages including phyloseq (https://doi.org/10.1371/journal.pone.0061217), microeco (https://doi.org/10.1093/femsec/fiaa255), and ggplot2. Taxa Erysipelotrichaceae and *Faecalibaculum rodentium* were chosen as the numerators for respective analyses based on statistical significance among groups.

### Fractionation and characterization of WPPC glycoconjugates

For the growth curve, RNA sequencing, and supernatant metabolomics experiments, a 30 kDa retentate fraction of the whey protein phospholipid concentrate (WPPC) was utilized. The procedure for obtaining this fraction was previously described.^40^ Characterization of the WPPC enriched in MFGM glycoconjugates (23%) compared to fluid bovine whole milk (1-2%) has also previously been described.^36,40^

### Bacteria and media

*Faecalibaculum rodentium* strain type strain ALO17 (DSM 103405) was obtained from the DSMZ-German Collection of Microorganisms and Cell Cultures (DSMZ) in lyophilized form and reconstituted according to instructions. Glycerol stocks (50% v/v) were stored at −80°C. Bacteria were routinely cultured in PYG medium (0.5% glucose) at 37°C in an anaerobic chamber (Coy Laboratory Products, Grass Lake, MI) with an atmosphere consisting of 5% carbon dioxide, 5% hydrogen, and 90% nitrogen. For all experiments, basal PYG (bPYG, no glucose) was used as base media.

### Growth curve in 30kD and constituent components

For each experiment, all work was performed in an anaerobic chamber. Briefly, glycerol stocks were streaked on to Columbia Blood Agar (CBA) plates with sheep blood (5%) (R01215, ThermoFisher Scientific) and allowed to grow until colonies formed. Individual colonies were picked and inoculated into PYG media for overnight growth. Overnight cultured strains were washed in bPYG three times and 2.5 μL of bacteria was inoculated into 160 μL of bPYG, or bPYG supplemented with 0.5% (wt/vol) of glucose, 40% (vol/vol) 30kD, or 1% (wt/vol) of lactose, galactose, mannose, N-acetlyglucosamine (GlcNAc), 2’-fucosyllactose (2’-FL), 3’-sialyllactose (3’-SL), 6’-sialyllactose (6’-SL), lacto-N-tetraose (LNT), and lacto-N-neotetraose (LnNT), fucose (Fuc), N-acetylneuraminic acid (Neu5Ac), and N-acetylgalactosamine (GalNAc). All carbon sources were obtained from MilliporeSigma, except for 2’-FL and 6’-SL (GenChem Co., South Korea). All media was filter sterilized with a 0.22 μm polyethersulfone (PES) membrane syringe filter (Genesee Scientific, San Diego, CA) before aliquoting into a 96 well plate. All wells were covered with 40 μL of mineral oil (MilliporeSigma) and incubated at 37°C in an anaerobic chamber (Coy Laboratory Products). Cell growth was monitored by assessing optical density (OD) 600 nm in real time using a BioTek PowerWave 340 plate reader (BioTek, Winooski, VT) every 30 minutes after 30 seconds of shaking at variable speed. For each condition, at least three technical replicates for three biological replicates were performed, and the average OD and standard error of the mean (SEM) were calculated to create growth curves over 24 hours.

### RNA extraction and transcriptomics

*F. rodentium* ALO17 from a single colony was grown in overnight culture, after which the cells were washed in bPYG and resuspended in bPYG to inoculate (10% vol/vol inoculum) into triplicates of 3 mL of PYG (0.5% glucose wt/vol) or triplicates of 40% (vol/vol) 30kDa retentate in bPYG media. Cells were grown until the late log/early stationary phase of growth (OD 0.51-0.67). Cells were harvested by 2,000 x g for 5 minutes of centrifugation at 4°C, treated with RNAlater Solution (Thermo Fisher^TM^ Scientific) according to the manufacturer’s protocol, and stored at −20°C until further use. Total RNA from the stored cells was isolated using the TRIzol® Reagent (Thermo Fisher Scientific) with modification, as previously published^40^ with the following modification: RNA elution was performed with 50 ul of RNase free water, and gDNA was removed by use of DNase (TURBO^TM^ DNase, Thermo Fisher Scientific).

Total RNA samples were sent to SeqCoast Genomics (Portsmouth, NH) for RNA-Seq analysis. RNA samples were prepared for sequencing using an Illumina Stranded Total RNA Prep Ligation Kit with Ribo-Zero Plus Microbiome (#20072063) with Illumina Unique Dual Indexes. Sequencing was performed on the Illumina NextSeq2000 platform using a 300-cycle flow cell kit to produce 2×150bp paired reads. 1-2% PhiX control was spiked into the run to support optimal base calling. Read demultiplexing, read trimming, and run analytics were performed using DRAGEN v4.2.7, an on-board analysis software on the NextSeq2000. Raw data was trimmed, aligned with the NCBI genome assembly ASM156445v1 (GenBank assembly GCA_00156445.1) and normalized by RPKM (reads per kilobase per million mapped reads) using CLC Genomics Workbench version Twelve (Qiagen), which was also used for differential gene expression (DEG) and statistical analysis. Gene functional categories were determined by literature search.

## Results

### Diet induced alteration of gut microbiota abundance in aging male Wistar rats

In our previous publication, we found that consumption of a high fat (HF) diet impaired cognitive function in aging male Wistar rats compared to a low fat (LF) diet (Sylvestre, et al. 2025, on bioRxiv). Interestingly, this impairment could be prevented by supplementation of the HF diet with 1.6% or 10% WPPC. To explore if the WPPC could be preventing cognitive impairment through the gut microbiota-brain axis, we first sought to determine if the diets altered the composition of the gut microbiome through 16S rRNA sequencing of rat feces. Beta diversity displayed as an NMDS plot (Figure 1A) depicts the largest differences between the LF and HF diet groups, with the 1.6% and the 10% WPPC supplementation increasingly more like the LF diet group.

**Figure 1.**
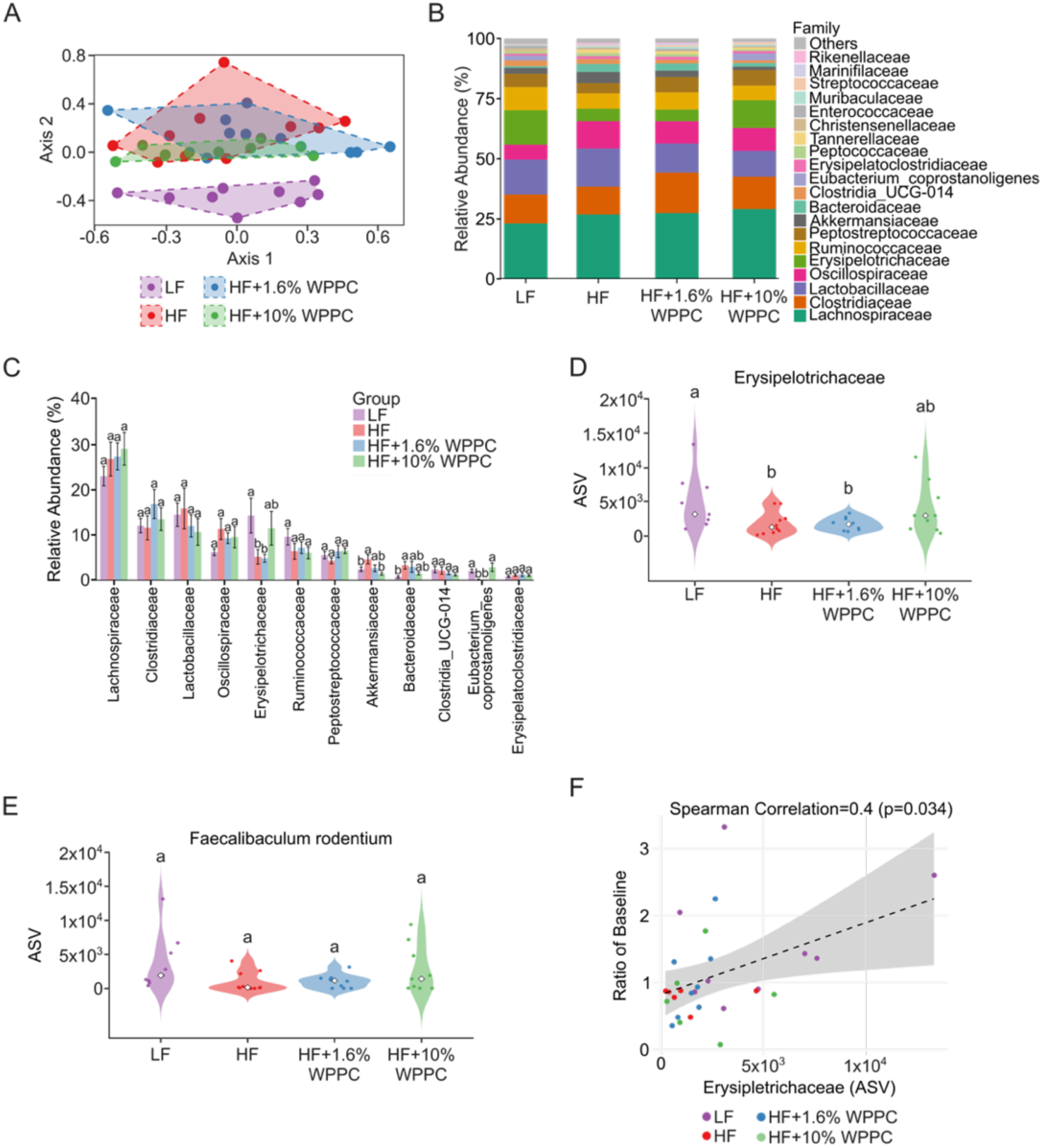
Microbiome composition analysis (16S rRNA sequencing) of Wistar rat fecal samples after 4 months on low fat (LF), high fat (HF), HF+1.6% WPPC, or HF+10% WPPC diets. (A) Non-Metric Multidimensional Scaling (NMDS) analysis based on Bray-Curtis dissimilarity showing microbiota clustering among different diet groups. (B) Relative abundance of the top 20 most abundant bacterial families across all experimental groups (taxa with a relative abundance of less than 1% were grouped as “Other”). (C) Bar plot of relative abundance of the top 12 most abundant families in all four experimental groups. (D, E) Violin plots depicting the abundance of *Erysipelotrichaceae* and *Faecalibaculum rodentium* across individual animals in each diet group. (F) Spearman correlation analysis between *Erysipelotrichaceae* abundance (ASV) and the baseline ratio, with a shaded area representing the 95% confidence interval. Statistical significance is indicated by different letters between experimental groups (p < 0.05, ANOVA with Duncan’s multiple range test).

Comparison of the abundance of the top 20 most abundant families across all four study groups indicates that *Lachnospiraceae, Clostridiaceae, Lactobacillaceae, Oscillospiraceae, and Erysipelotrichaceae* are the top five most abundant families present across all study groups, collectively accounting for greater than 60% of all families’ relative abundance (R.A.) (Figure 1B). Closer examination of the top twelve most abundant families’ abundance across all four study groups indicated that one family, *Erysipelotrichaceae,* had a pattern of relative abundance that appeared to mirror cognitive performance patterns in the rats: high R.A. in the LF diet (14.3%), significantly reduced R.A. in the HF diet (5.18%, P=0.015), and elevated R.A. in the HF+10% WPPC diet (11.5%, P=0.044) (Figure 1C). This pattern of abundance was not observed for any of the other families identified by 16S rRNA sequencing. Closer examination of the abundance of the *Erysipelotrichaceae* for each of the individual rats per study group demonstrated a wide range of abundance for the LF (14.3 ± 4.0) and HF+10% WPPC groups (11.5 ± 3.8), but reduced variation in abundance for the HF (5.2 ± 1.6) and HF+1.6% WPPC groups (4.8 ± 0.8) (Figure 1D). Within the family *Erysipelotrichaceae,* the species *Faecalibaculum rodentium* (*F. rodentium*) was identified and its abundance measured across all four study groups (Figure 1E), which mirrored the family level abundance patterns. The differences in *F. rodentium* abundance were numerically higher in the LF and HF+10% WPPC groups compared to the HF and HF+1.6%WPPC groups, but none of the differences were statistically significant (LF vs HF, P=0.079; HF vs HF+10% WPPC, P=0.140).

### Positive correlation between Erysipelotrichaceae family abundance and hippocampal memory storage

To determine if there were any correlations between the abundance of *Erysipelotrichaceae* or *Faecalibaculum rodentium* taxa with cognitive function in the rats, we performed correlation analysis between the family and species level taxa abundance with the discrimination index, effective 50% dose, and ratio of baseline measured in the rats. Briefly, each measurement provides different information on memory or cognitive function. Discrimination index measures how well an animal can distinguish between two items. Effective 50% Dose measures how much of a treatment is needed to see a meaningful improvement in cognitive function, while ratio of baseline compares performance after an intervention and reflects memory storage.

Comparison of the abundance of *Erysipelotrichaceae* taxa (Spearman correlation=0.25, P=0.125) and *Faecalibaculum rodentium* (Spearman correlation=0.29, P=0.081) with the discrimination index did not indicate any significant correlations (Supplemental Figure 1A and B, respectively). Likewise, comparison of the abundance of *Erysipelotrichaceae* taxa (Spearman correlation=0.17, P=0.387) and *Faecalibaculum rodentium* (Spearman correlation=0.24, P=0.203) with the Effective 50% dose did not indicate any significant correlations (Supplemental Figure 1C and D, respectively). However, while the *Faecalibaculum rodentium* correlation with the ratio of baseline was not significant (Spearman correlation=0.28, P=0.14) (Supplemental Figure 1E), the *Erysipelotrichaceae* taxa did positively and significantly correlate with the ratio of baseline (Spearman correlation=0.4, P=0.034) (Figure 1F). This suggests a potential relationship between the abundance of the *Erysipelotrichaceae* taxa and the ability for memory storage in the rats.

### Faecalibaculum rodentium strain ALO17 growth on 30kD and its constituents

To determine if *Faecalibaculum rodentium* could grow in isolation on WPPC glycoconjugates, we obtained the type strain ALO17 for growth curve analysis. Previously, we demonstrated the enrichment of milk fat globule membrane (MFGM) glycoconjugates from the WPPC in a 30kDa retentate fraction, and that it could be used as a sole carbon source to study bacteria growth.^40^ Using basal PYG media as a negative control (Figure 2A) and glucose as a positive control (Figure 2B), we found that ALO17 grew on the 30kDa retentate over 24 hours (Figure 2C).

**Figure 2.**
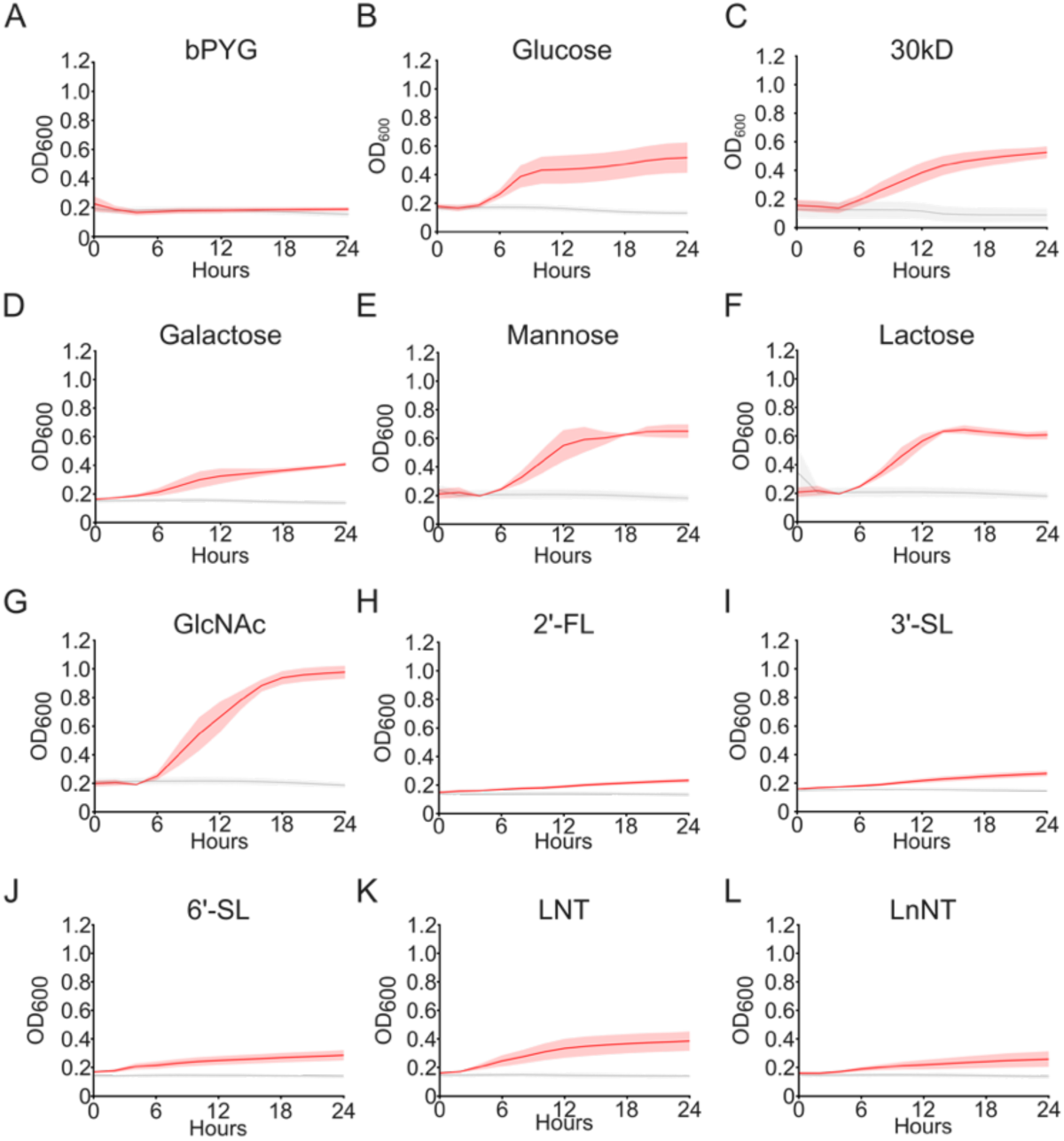
Growth of *Faecalibaculum rodentium* strain ALO17 on different WPPC-constituent substrates. Growth curves are of media (grey) and *Faecalibaculum rodentium* strain ALO17 (red) over a 24 hour period. Media blanks and with ALO17 inoculum were grown in the following conditions: (A) bPYG, (B) glucose, (C) 30kD fraction, (D) galactose, (E) mannose, (F) lactose, (G) N-acetylglucosamine (GlcNAc), (H) 2’-fucosyllactose (2’-FL), (I) 3’-siallyactose (3’-SL), (J) 6’-siallyactose (6’-SL), (K) lacto-N-tetraose (LNT), and (L) lacto-N-neotetraose (LnNT). Each experiment was repeated at least three times.

Previous analysis indicated that glycan substrates present in the WPPC enriched 30kDa include glucose, galactose, lactose, mannose, and N-acetylglucosamine (GlcNAc).^40^ As such, we tested the growth of ALO17 on each of these constituent components and found growth (Figure 1D-G). We further tested ALO17’s ability to grow on other milk components but found that it did not grow on 2’-FL, 3’-SL, 6’-SL, LNT, or LnNT (Figure 1H-L), or GalNAc, Fucose, or Neu5Ac (Table 1). Of all substrates, ALO17 appeared to demonstrate the most robust growth on GlcNAc (Figure 2G).

**Table 1.**

| Sample | No<br>carbohydrates | 30kD | 30kD components |  |  |  | Man | GalNAc | Fuc | 2'-FL | Neu5Ac | 3'-SL | 6'-SL | LNT | LnNT |
| --- | --- | --- | --- | --- | --- | --- | --- | --- | --- | --- | --- | --- | --- | --- | --- |
|  |  |  | Glc | Gal | Lac | GlcNAc |  |  |  |  |  |  |  |  |  |
| ALO17 | - | ++ | ++ | + | ++ | +++ | ++ | - | - | + | - | + | + | + | + |
| Mean Maximum<br>OD600 by 24 hours | 0.19 | 0.53 | 0.52 | 0.41 | 0.65 | 0.98 | 0.65 | 0.184111 | 0.201778 | 0.23 | 0.176222 | 0.267 | 0.29 | 0.39 | 0.26 |
| SEM Maximum OD600<br>by 24 hours | 0.01 | 0.04 | 0.11 | 0.01 | 0.03048 | 0.045671 | 0.047914 | 0.020113 | 0.023898 | 0.014987 | 0.01612 | 0.019062 | 0.035741 | 0.068315 | 0.055205 |
| Media mean | 0.16 | 0.1 | 0.14 | 0.14 | 0.19 | 0.19 | 0.19 | 0.18 | 0.17 | 0.13 | 0.15 | 0.15 | 0.14 | 0.14 | 0.14 |
| Media baseline range<br>(min-max) | 0.05-0.31 | 0.01-0.20 | 0.03-0.23 | 0.12-0.17 | 0.14-0.28 | 0.14-0.27 | 0.14-0.28 | 0.14-0.28 | 0.13-0.32 | 0.11-0.15 | 0.12-0.26 | 0.13-0.16 | 0.12-0.16 | 0.12-0.18 | 0.12-0.18 |

### Transcriptomic analysis of Faecalibaculum rodentium ALO17 during growth on 30kDa retentate compared to glucose

To determine in growth on the MFGM enriched 30kDa retentate induced changes in gene expression compared to glucose, we performed RNA sequencing on the type strain ALO17 grown on 0.5% glucose or 40% 30kD fraction. The different substrate concentrations were used due to the different growth inducing capacities of glucose (a monomer) compared to the 30kD retentate (a complex MFGM glycoconjugate-enriched substrate).^40^ The PCA plot of the transcriptomic data indicates a clear distinction in gene expression between the glucose and 30kD groups (Figure 3A). In total 2,887 genes were identified in the ALO17 genome (Figure 3B), 961 of which were differentially expressed (FDR<0.05) between the two groups (33% of all genes). Of the 961 differentially expressed genes (DEGs), 547 genes (19% of all genes) were upregulated by 30kD consumption, and 414 genes (14% of all genes) were upregulated by glucose consumption.

**Figure 3.**
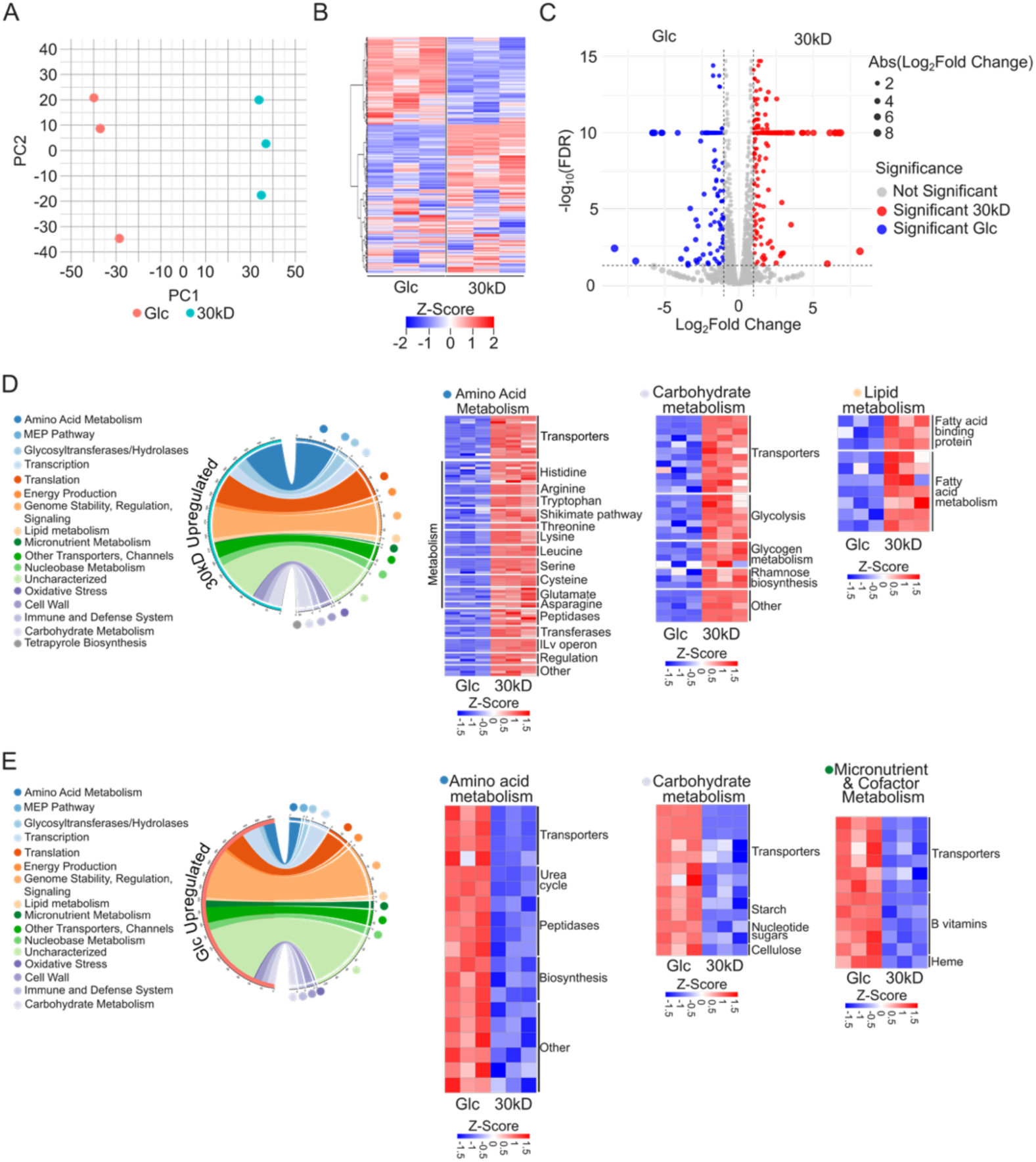
Transcriptomic analysis of *Faecalibaculum rodentium* ALO17 during growth on 30kDa retentate or glucose. (A) PCA plot of transcriptomic data showing distinct gene expression profiles between glucose (red) and 30kDa retentate (blue) conditions. (B) Heatmap depicting the expression of the ALO17 genome (2,887 genes) with 961 differentially expressed genes (DEGs, FDR<0.05), 547 upregulated by 30kDa and 414 by glucose. (C) Volcano plot displaying DEGs with |log2FoldChange| > 1 (206 genes upregulated on 30kDa; 114 on glucose). (D) Functional classification of 30kDa upregulated genes across 17 biological categories, highlighting pathways involved in amino acid, carbohydrate, and lipid metabolism, with heatmaps showing genes linked to these processes. (E) Functional classification of glucose upregulated genes, highlighting pathways involved in amino acid, carbohydrate, and micronutrient and cofactor metabolism, with heatmaps showing genes linked to these processes. Statistical significance determined by FDR<0.05.

A total of 320 genes had FDR<0.05 and |log2FoldChange| > 1, as depicted in the volcano plot (Figure 3C). Of these, 206 genes were upregulated on 30kD (red circles) and 114 genes were upregulated on glucose (blue circles). To understand the biological context of the genes upregulated by the 30kD (Figure 3D) or glucose (Figure 3E), a literature search was performed on all annotated genes, resulting in manually curated classification into seventeen biological categories. Classification of the 547 30kD upregulated genes was distributed as follows (Figure 3D): amino acid metabolism (88), methylerythritol phosphate (MEP) pathway (3), glycosyltransferases/hydrolases (23), transcription (28), translation (68), energy production (10), genome stability, regulation, signaling (73), lipid metabolism (10), micronutrient metabolism (3), other transporters, channels (34), nucleobase metabolism (19), uncharacterized (98), oxidative stress (2), cell wall (23), immune and defense system (13), carbohydrate metabolism (31), tetrapyrrole biosynthesis (1).

Of the 30kD upregulated genes, those involved in amino acid metabolism (88), carbohydrate metabolism (31), and lipid metabolism (10) are highlighted as heatmaps (Figure 3D) and were further analyzed due to their potential biological relevance in preventing cognitive impairment (i.e., microbial metabolite secretion). For amino acid metabolism 88 DEGs were identified, of which 16 genes encode for transporters and 72 genes encode for enzymes involved in protein and amino acid metabolism. Of the transporters, all 16 appear to be involved in amino acid and peptide import, reflecting the protein rich content of the 30kD fraction. The remaining 72 DEGs are involved in a variety of metabolic processes, including for the metabolism of histidine (8), arginine (4), tryptophan (4), threonine (2), lysine (5), leucine (4), serine (6), cysteine (4), glutamate (5), asparagine (2), along with the Shikimate pathway (5) which produces aromatic amino acids. In addition, 30kD consumption upregulated the expression of peptidases (6), transferases (4), and genes in the ILv operon (5), regulation of amino acid metabolism (4), and others (4).

In addition to amino acid metabolism, 31 genes involved in carbohydrate metabolism were significantly upregulated by 30kD consumption (Figure 3D). Of these, 12 are for transporters, 19 are involved in metabolic processes including glycolysis (7), glycogen metabolism (4), rhamnose biosynthesis (3), and other processes (5). Of the transporters, the majority are annotated as phosphotransferase system (PTS) transporters for fructose, mannose, glucose, lactose, or cellobiose. For lipid metabolism, upregulated genes are annotated as being fatty acid binding proteins (3) or involved in fatty acid metabolism (7), the latter of which are involved both fatty acid catabolism and biosynthesis.

Classification of the 414 glucose upregulated genes was distributed as follows (Figure 3E): amino acid metabolism (19), methylerythritol phosphate (MEP) pathway (2), glycosyltransferases/hydrolases (7), transcription (30), translation (35), energy production (5), genome stability, regulation, signaling (85), lipid metabolism (5), micronutrient metabolism (12), other transporters, channels (26), nucleobase metabolism (13), uncharacterized (111), oxidative stress (2), cell wall (12), immune and defense system (10), and carbohydrate metabolism (13).

For amino acid metabolism 19 DEGs were identified, of which 4 genes encode for transporters and 15 genes encode for enzymes involved in protein and amino acid metabolism. In contrast to the 30kD upregulated amino acid transporters, the four glucose upregulated transports appear to be involved in amino acid export, particularly for threonine and serine. The remaining 15 DEGs are involved in several metabolic processes, including the urea cycle (2), amino acid biosynthesis (3), peptidases (4), and others (3). For carbohydrate metabolism, 13 genes were significantly upregulated by glucose consumption (Figure 3E). Of these, 8 are for transporters, 5 are involved in metabolic processes including starch metabolism (2), nucleotide sugar metabolism (2), and cellulose metabolism (1). Of the transporters, the majority are annotated as phosphotransferase system (PTS) transporters for lactose or cellobiose, or ABC transporter permeases.

For the glucose condition, 12 genes involved in micronutrient and cofactor metabolism were significantly upregulated. Of these, the majority were annotated as transporters (6), followed by those involved in B vitamin metabolism (5) and heme biosynthesis (1). Of the transporters, one is involved in folate (vitamin B9), four in mineral transport, and one in L-ascorbate (vitamin C) transport. For the five B vitamin metabolism genes, two are involved in folate (vitamin B9) metabolism, one in thiamine (vitamin B1) biosynthesis, one in cobalamin (vitamin B12) biosynthesis, and one in the metabolism of pyridoxine (vitamin B6).

### 30kD fraction stimulated Faecalibaculum rodentium production of histidine

Of the 30kD upregulated DEGs, the histidine biosynthesis pathway appeared to be strongly stimulated by 30kD consumption (Figure 4A). Eight genes involved in histidine biosynthesis were identified, including: HisA, HisF, HisH, HisE, HisD, HisG, aalo17_RS07600, and aalo17_RS11820.

**Figure 4.**
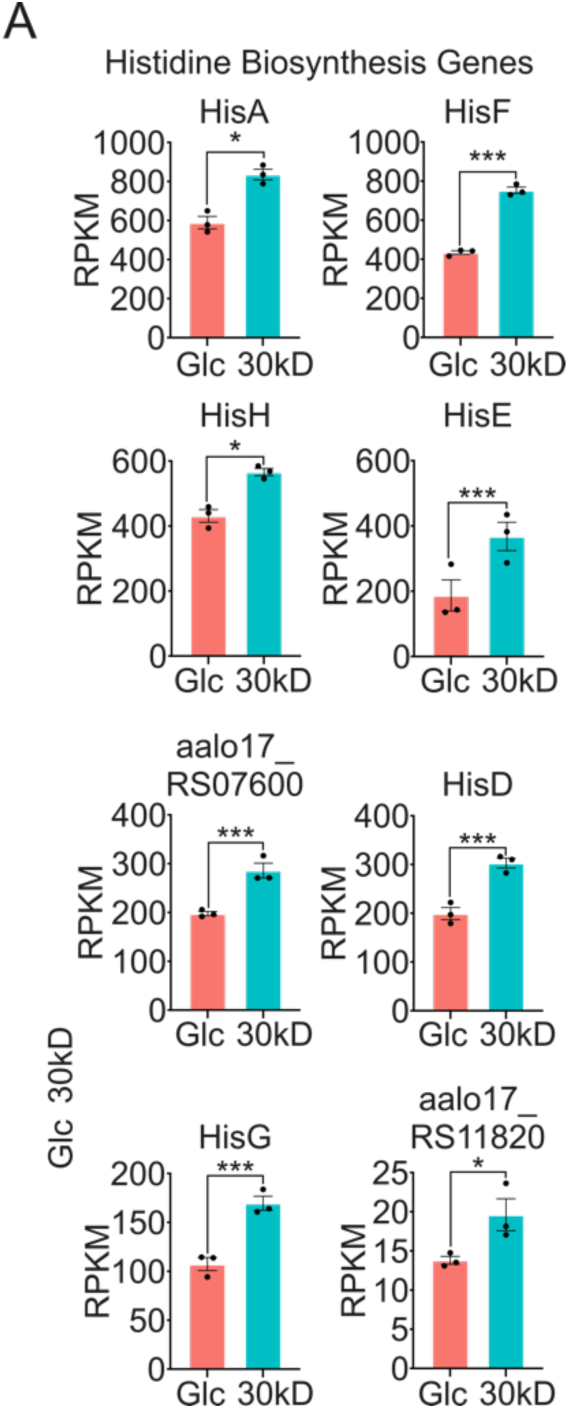
30kD consumption stimulated *F. rodentium* histidine production and secretion. (A) Consumption of 30kD significantly upregulated genes involved in histidine biosynthesis compared to consumption of glucose (FDR<0.05).

## Discussion

### MFGM-enriched WPPC as a potential dietary intervention to prevent cognitive impairment

The MFGM can exert beneficial effects in the host through microbe dependent and independent mechanisms.^41–43^ Limited studies have examined changes in the gut microbiota in concert with changes in cognitive function and/or the effect of MFGM supplementation.^7,9–11^ O’Mahony, et al. found that provision of MFGM to adult mice significantly increased the abundance of the family Erysipeltrichaceae compared to mice not receiving MFGM, which was the greatest effect seen at the family level.^11^ Wu, et al., found that prophylactic MFGM treatment enriched Faecalibaculum abundance in feces in mice treated with DSS, compared to mice treated with DSS alone.^7^ Berding found that supplementation of MFGM (along with prebiotics) to infant formula increased the fecal abundance of an Erysipeltrichaceae genus compared to piglets not receiving the supplementation.^10^

Commercial sources of MFGM are either whey or cream derived and have been found to vary in protein and lipid composition.^35–39^ By leveraging an abundant MFGM enriched dairy co-product, the WPPC, this study contributes to our understanding of how the MFGM modulates gut microbiota composition and function in association with improved cognitive function in aging male Wistar rats fed a HF diet. The goal of this study was to explore if a potential nutrient-gut microbe-brain axis mechanism exists for how the WPPC prevents cognitive impairment, which could present a novel avenue for the WPPC as a dietary intervention to prevent neurodegeneration and cognitive impairment. To date, pharmacological approaches dominate treatments for dementia, with two controversial drugs approved by the FDA for AD treatment in late-stage cases. Non-pharmacological prevention and treatment modalities that target the gut microbiota represent novel avenues for reducing chronic disease. In agreement with our research, others have found that provision of MFGM can mitigate cognitive impairment in adult mice,^15^ and have other beneficial effects in aging such as suppressing the progression of age-related alterations in neuromuscular junctions (when combined with exercise).^44^ A recent randomized controlled trial of 107 older adults (mean age 62.94 years old) found that supplementation of a MFGM enriched whey protein powder improved cognitive function over a 12-month period, although the results were not statistically significant.^45^ As such, there is interest and potential for the use of MFGM-enriched fractions to prevent cognitive impairment in humans.

### Erysipelotrichaceae and Faecalibaculum rodentium abundance is influenced by diet and consumption of the WPPC

In this study, we identified that the family *Erysipelotrichaceae* and specifically the species *Faecalibaculum rodentium* were abundant on the LF diet, reduced by the HF diet, and restored to abundance by HF+10%WPPC supplementation (Figure 1C-E). As such, we chose to examine this bacteria species for a potential mediator for WPPC consumption preventing cognitive impairment.

The bacteria species *Faecalibaculum rodentium* (*F. rodentium*) was first isolated in 2015 from the feces of a nine-month-old female C57BL/6 J laboratory mouse fed a standard experimental diet by Chang, et al. in the Republic of Korea, with the type strain designated as ALO17.^46^ The species name comes from faecalis pertaining to feces, baculum meaning small rod, and the Latin rodere, which means “to gnaw,” indicating a small rod baculum isolated from rodent species. The anaerobic, gram positive, rod-shaped, non-spore forming, non-motile bacterium (BSL1) was originally isolated under anaerobic conditions using DSM 104 medium. TEM images indicate it arranges in pairs or chains measuring at 0.9-1.2 μm in width and 1.5-2 μm in length. The main product of its glucose fermentation is lactic acid with small amounts of ethanol, acetic acid and butyric acid also produced. *F. rodentium* ALO17 has a single circular DNA chromosome with a genome size of 2.54Mb, with an estimated 2583 ORFs and a GC content of 54% with no plasmids.^47^

*F. rodentium* is a member of the class Erysipelotrichia and phylum firmicutes, which are found in the gastrointestinal tracts of diverse species including rodents,^48^ swine,^49^ chickens,^50^ and the rhinoceros beetle.^51^ The family *Erysipelotrichaceae* is also present in humans,^52,53^ and to date 10 genera have been identified in humans.^54^ In mice, the abundance of the *Erysipelotrichaceae* family was found to be most abundant in middle adult aged mice (589 days old) compared to young adult (174 days old) and old adult mice (857 days old).^55^ In early life, the genus Faecalibaculum increases in abundance from 4, 8 to 12 weeks of age in wild type C57BL/6 mice.^56^ *F. rodentium* was found to be decreased in abundance when FMT was collected at ZT23 (lights off) compared to ZT11 (lights on), indicating abundance may be affected by circadian rhythms.^57^ In agreement with this study, it was previously found that wild-type mice consuming normal chow (5.2% fat) had much higher abundance of *Erysipelotrichaceae* compared to mice placed on a high-fat diet (34.9% fat).^58^ Sun, et al. (2019) found that fecal microbiota transplant (FMT) from wild type mice could improve cognitive defects in APPswe/PS1dE9 transgenic mice, which was associated with an increase in abundance in the family *Erysipelotrichaceae* and the genus Faecalibaculum.^59^

In humans, the taxon Erysipelotrichaceae UCG-003 is more abundant in healthy old age than in non-healthy old age,^60^ while cc115 (family *Erysipelotrichaceae*) is more abundant in healthy controls than patients with Alzheimer’s disease.^61^ Conversely, in people living with HIV the relative abundance of Erysipelotrichaceae UCG-003 was inversely correlated with age acceleration, suggesting that its increased abundance is associated with slower rates of biological aging.^62^ Collectively, it appears that the family *Erysipelotrichaceae* and species *F. rodentium* are associated with healthy aging and improved cognitive function across species, suggesting a potential conserved gut-brain axis mechanism for promoting brain health.

### Consumption of the 30kD fraction shifts the metabolic activity of F. rodentium relative to glucose

Consumption of the 30kD compared to glucose induced broad differences in gene expression in the *F. rodentium* strain ALO17 (Figure 3). Notably, consumption of 30kD upregulated 88 genes involved in amino acid metabolism, while consumption of glucose only upregulated 19 genes involved in amino acid metabolism. Likewise, compared to glucose the 30kD significantly upregulated more genes classified as glycosyltransferases/hydrolases (23 vs 7 genes), and genes involved in translation (68 vs 35 genes), carbohydrate metabolism (31 vs 13 genes), energy production (10 vs 5 genes), lipid metabolism (10 vs 5 genes), and cell wall (23 vs 12 genes). Notably, the 16 animo acid transporters upregulated by 30kD are all involved in import, while the four amino acid transporters upregulated by glucose appear to be involved in amino acid export, particularly for threonine and serine. This may reflect the high protein content of the 30kD compared to the glucose media, which drove ALO17 growth on amino acid fermentation (import) compared to glucose fermentation.

In contrast, consumption of glucose upregulated more genes involved in micronutrient and cofactor metabolism compared to the 30kD (12 vs 3 genes). The three 30kD micronutrient related genes are involved in mineral acquisition or metabolism, while the glucose upregulated genes appear to indicate a concerted production in vitamins B1, B6, B9, and B12. As *F. rodentium* has been demonstrated to produce B vitamins when grown on glucose, this finding agrees with previous studies.^48^

One interesting gene that was significantly upregulated by glucose is choloylglycine hydrolase, also known as bile salt hydrolase (BSH). BSH hydrolyzes conjugated bile salts into deconjugated bile acids (Bas), which are crucial signaling molecules in multiple metabolic processes such as lipid and glucose metabolism. While the function of this gene being activated by glucose consumption is not clear, it indicates a role for how MFGM may regulate how *F. rodentium* responds to surrounding bile acids in the gut.

### 30kD fraction stimulated F. rodentium production of histidine

L-histidine is an essential amino acid, meaning that it must be consumed in the diet or produced but gut microbiota. The unique biochemical properties and physiological functions of histidine have resulted in its use as a nutritional supplement to treat a plethora of conditions including rheumatoid arthritis, anemia, fatigue, inflammatory bowel diseases (IBD), ocular diseases, and neurological diseases.^63^ Histidine is mainly obtained in the diet from proteins; the WHO/FAO recommended dietary requirement is 8-12 mg/kg/day for adults 19 years and older (<1.5g histidine/day), however daily intake has been reported to range between 1.52g/day (vegans) to 2.12 g/day (meat-eaters).^64^

Histidine is one of the least abundant amino acids in humans and of all organs is most abundant in skeletal muscle.^65^ A deficiency in histidine has been implicated in several disease contexts in multiple species. In largemouth bass (*Micropterus salmoides*), histidine deficiency suppressed the activity of intestinal antioxidant enzymes and induced intestinal endoplasmic-reticulum stress and proinflammatory cytokine production.^66^ Histidine has been found to prevent the pathophysiology of colitis in rodents^67^ and is reduced in the serum of IBD patients.^68^ In rodents, histidine supplementation ameliorated compromised memory function as induced by sleep deprivation.^69^ In humans with sleep disruption and high fatigue, histidine was able to significantly reduce fatigue and improve working memory.^70^

Histidine is also important as a precursor of histamine. Histidine consumed in the diet or produced by gut microbes can enter systemic circulation through neutral amino acid transporters^71^ and is able to cross the blood brain barrier (BBB) and enter the brain.^72^ Throughout the brain but most pronounced in the posterior hypothalamus region, histidine is converted to histamine by the enzyme histidine decarboxylase (HDC) which is expressed in histaminergic neurons.^73^ As a neurotransmitter, histamine controls multiple functions such as memory, appetite, the sleep-wake cycle, and stress response.^74^ Histamine has been found to play a role in short-term recognition memory in rats,^75^while the use of H_3_ receptor (H_3_R) antagonists (which increase histamine release in the brain) improved wild type mice memory, but not histamine-deficient mice memory, in the object recognition test.^76^ In models of cognitive disorders, use of H_3_R antagonists improved object recognition in mice that received amyloid peptide injections,^77^ while long-term treatment with the same H_3_R antagonist prevented cognitive deficits in a tau transgenic mouse model.^78^ Interestingly, insufficient intake of histidine has been found to reduce brain histamine abundance and cause anxiety-like behaviors in male mice,^79^ while HDC knock out mice exhibit locomotor behaviors the mimic Tourette’s syndrome.^80^ In infants, elevated histidine levels were associated with composite cognition above versus below median values,^53^ while fecal *Erysipelotrichaceae* abundance was positively correlated with fecal histidine levels. Notably, quantification of histamine in nine *postmortem* Alzheimer brains revealed a reduction in histamine content in the temporal cortex (53% of the control value), hippocampus (43%), and the hypothalamus (42%) compared to five age-matched controls.^81^

In this study, we identified that *Faecalibaculum rodentium* is able to grow on the 30kD WPPC fraction, and that growth on the 30kD significantly increased the expression of genes associated with histidine biosynthesis compared to growth on glucose (Figure 4A). In agreement with our finding, Zeng, et al. found that *F. rodentium* can produce and secrete histidine, which protected against colitis in *Il10−/−* mice.^48^ The authors identified a suite of histidine-associated genes in *F. rodentium* which were upregulated in expression by glucose in the media compared to glucose deficient media. In addition, Zeng, et al. analyzed metagenomic datasets from three human cohorts and found that most histidine biosynthesis genes were upregulated in non-IBD individuals compared to patients with CD.

In other contexts, *F. rodentium* has been observed to exert diverse beneficial effects. Zagato, et al. (2020) found *F. rodentium* is underrepresented during early tumorigenesis and can prevent tumor growth in two mouse models through its metabolic products, especially butyrate which was found to reduce tumor cell proliferation.^56^ Interestingly, the authors also found that, compared to healthy individuals, patients with large adenomas have a reduction in the family *Erysipelotrichaceae,* especially the species *Holdemanella biformis.* In a separate study, Cao, et al. (2022) found that *F. rodentium* can regulate duodenal epithelial turnover through remodeling gut luminal retinoic acid signaling, which can block immunological eosinophils from curbing epithelial turnover.^82^ As aging is marked by a decline in the regenerative potential of intestinal stem cells, and thus reduced rates of intestinal epithelial cell turnover,^83,84^ this suggests an immune system mediated mechanism for how *F. rodentium* may be able to promote healthy aging by maintaining homeostatic rates of turnover in the gut.

Still, others have found that *F. rodentium* is associated with negative health outcomes such as its ability to displace commensal Th17-inducing microbiota in mice on high sugar diets, thereby depleting Th17 cells that protect against metabolic syndrome through regulating lipid absorption.^85^ Interestingly, Wang, et al. found that ingestion of *F. rodentium* in *Ephx2* knock out mice caused depression-like phenotypes, indicating genetic contexts in which this species could exert negative effects.^86^

Limitations to this study include the lack of functional validation of histidine secretion by *F. rodentium* as a mechanism to prevent HF-diet induced cognitive impairment in aging male Wistar rats. In this paper, we present a rationale for future work to test this mechanism as a means by which the WPPC prevents cognitive impairment as induced by the HF diet. However, at present our data remains largely correlative in nature. In addition, it would be interesting to identify what specific components of the WPPC and the 30kD are responsible for upregulating the histidine biosynthesis genes in *F. rodentium,* and to understand how histidine is preventing cognitive impairment in the brain. Such questions would be valuable to explore in future research.

## Abbreviations List

WPPC: Whey Protein Phospholipid Concentrate
MFGM: Milk Fat Globule Membrane
LF: Low Fat
HF: High Fat
HF1.6%: High Fat + 1.6% Whey Protein Phospholipid Concentrate
HF10%: High Fat + 10% Whey Protein Phospholipid Concentrate

## Data Availability

The RNA sequencing data presented in the study are deposited in the NCBI SRA database repository, BioProjectID: PRJNA1235763.

## Acknowledgements

The authors would like to thank Gallo for the donation of WPPC which was used to generate the 30kD fraction. We also thank Audrey Pimienta for assistance in laboratory work.

## Funding Acknowledgements

This research was supported by funds from the Shields Endowed Chair in dairy Food Science (DAM), the National Dairy Council (DB), and California Dairy Research Foundation (DB). Rat feces was obtained from animal subjects that received approval by the Institutional Animal Care and Use Committee (IRB approval #22543). The authors do not have any conflicts of interest.

---

**Supplemental Figure 1.**
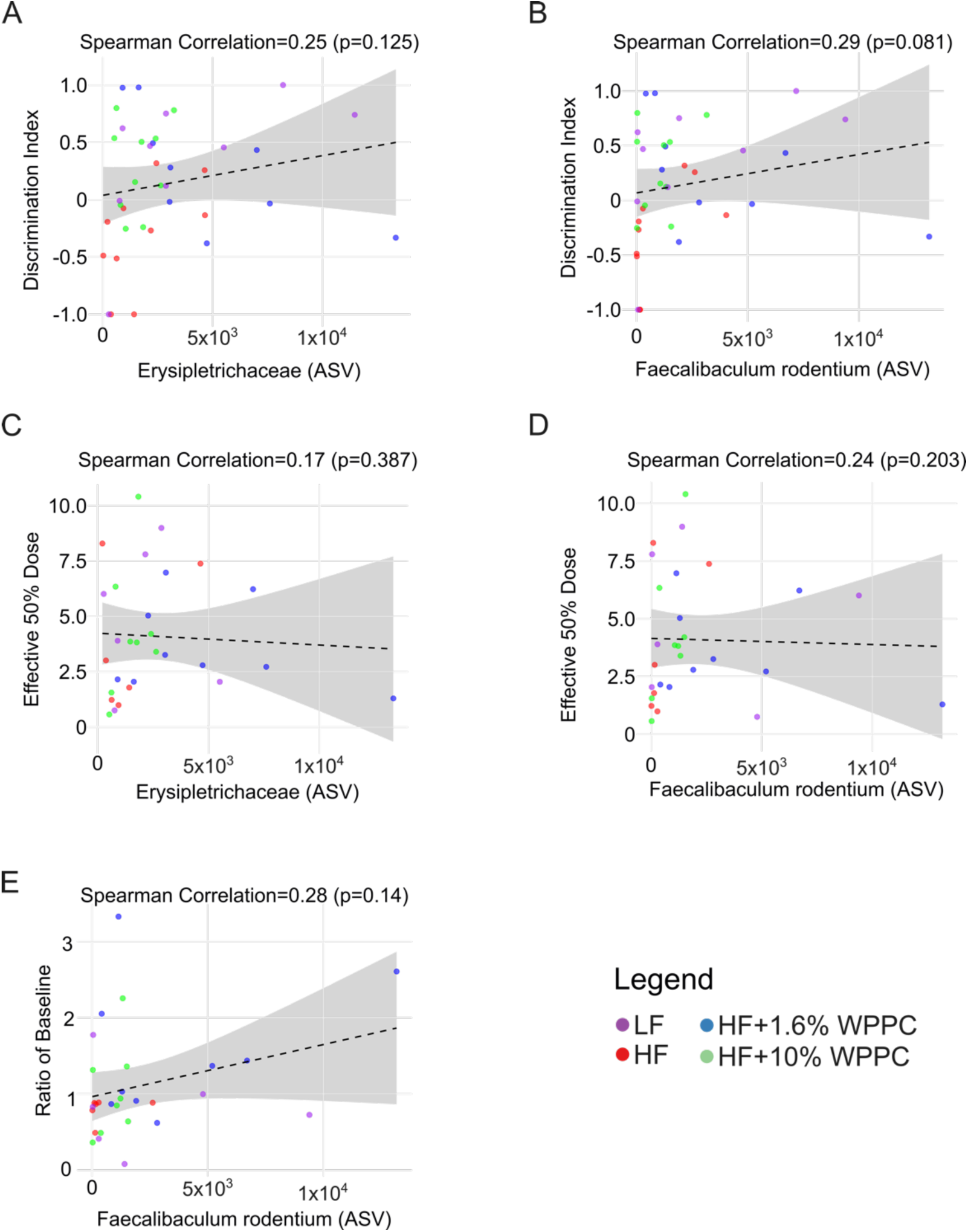
Correlation of *Erysipelotrichaceae* and *Faecalibaculum rodentium* abundance with behavioral metrics. (A, B) Spearman correlation analysis between the abundance of *Erysipelotrichaceae* (ρ = 0.25, *P* = 0.125) and *Faecalibaculum rodentium* (ρ = 0.29, *P* = 0.081) with the discrimination index, showing no significant correlations. (C, D) Correlation analysis with the Effective 50% dose for *Erysipelotrichaceae* (ρ = 0.17, *P* = 0.387) and *Faecalibaculum rodentium* (ρ = 0.24, *P* = 0.203), also showing no significant associations. (E) Correlation of *Faecalibaculum rodentium* abundance with the ratio of baseline (ρ = 0.28, *P* = 0.14), which was not significant. Statistical significance was set at *P* < 0.05.

